# An oviduct glycan increases sperm lifespan by diminishing ubiquinone and production of reactive oxygen species

**DOI:** 10.1101/2023.01.08.523174

**Authors:** Jennifer R. Hughes, Katie J. McMorrow, Nicolai Bovin, David. J. Miller

## Abstract

Sperm storage by females after mating for species-dependent periods is used widely among animals with internal fertilization to allow asynchrony between mating and ovulation. Many mammals store sperm in the lower oviduct where specific glycans on epithelial cells retain sperm to form a reservoir. Binding to oviduct cells suppresses sperm intracellular Ca^2+^ and increases sperm longevity. We investigated the mechanisms by which a specific oviduct glycan, 3-O-sulfated Lewis X trisaccharide (suLe^X^), prolongs the lifespan of porcine sperm. Using targeted metabolomics, we report that binding to suLe^X^ diminishes the abundance of the precursor to ubiquinone and suppresses formation of fumarate, a specific citric acid cycle component, diminishing the activity of the electron transport chain and reducing the production of harmful reactive oxygen species (ROS). The enhanced sperm lifespan in the oviduct may be due to suppressed ROS production as many reports have demonstrated toxic effects of high ROS concentrations on sperm.

## Introduction

Storage of sperm in the female reproductive tract allows for successful fertilization when ovulation and mating are not synchronized tightly. The ability of the female tract to store sperm is nearly ubiquitously conserved among animals with internal fertilization including a wide variety of reptiles, fish, birds,[1] insects, and mammals[2-4]. The reservoir location sometimes differs between species[1, 4, 5], implying repeated evolution of sperm storage systems. Insects, salamanders, and newts store sperm in specialized sacs[6]. Birds and reptiles store sperm in dead-end tubules that are found in the uterovaginal junction[4, 5]. In contrast, mammals store sperm primarily in folds of the lower oviduct, the isthmus[5, 7, 8].

Some animals, such as species of ants and bees, store sperm for the lifetime of the female[4, 9]. Among mammals, it is notable that some species of bats store sperm for months[10] although sperm storage in most mammals is for shorter periods, usually days[7]. Sperm binding and retention within the mammalian isthmus favors live, motile, and morphologically normal sperm[3]. Sperm that have not completed capacitation, the final maturation before fertilizing oocytes, also are preferentially bound by the isthmus[3]. Additionally, sperm binding to the oviduct reduces the frequency of polyspermy[11]. These discoveries led to the hypotheses that the functional sperm reservoir in mammals suppresses capacitation, selects functional sperm, and provides a controlled release of sperm for fertilization[12].

In mammals studied to date, sperm are stored in the oviduct by binding to glycans on the epithelial cell surface[13-16]. Porcine sperm bind preferentially to 6-sialylated multi-antennary oligosaccharides and Lewis X and 3-O-sulfated Lewis X (suLe^X^) trisaccharides with high affinity as demonstrated by glycan array[15, 17]. The surface epithelium within the porcine isthmus maintains a high abundance of glycoproteins and glycolipids decorated with 6-sialylated structures and Lewis X trisaccharides[14]. The Lewis X trisaccharide is composed of an N-acetyl-glucosamine residue with a fucose residue attached in an α3 linkage and galactose residue attached in a 4 linkage[18]. Both the 6-sialylated and Lewis X structures are components of glycans that are attached in glycoproteins primarily to asparagine residues[14]. Binding to suLe^X^ *in vitro* mimics an important function of the isthmus. When coupled to beads, suLe^X^ enhanced the viability of bound sperm over 24 hr[19]. These changes in sperm viability, observed under *in vitro* capacitating conditions, indicate that binding to immobilized suLe^X^ induces changes in sperm function.

Sperm are transcriptionally silent cells with a very limited amount of mRNA and ribosomes. They rely on environmental factors for signals to begin their final maturation, known as capacitation, including hyperactivation and acrosomal exocytosis[20], leading to fertilization of an oocyte[21]. Delaying sperm progress through the terminal differentiation process of capacitation may be critical to maintaining viable sperm within the female reproductive tract.

Capacitation includes multiple events that are interconnected and complex. These processes include loss of membrane cholesterol, increased intracellular Ca^2+^, increased tyrosine phosphorylation of some proteins, and an altered (hyperactivated) motility pattern[20, 22]. Previous research has shown that increases in intracellular Ca^2+^ are suppressed in sperm bound to oviduct glycans, including suLe^X^, but no such change in either motility pattern or protein tyrosine phosphorylation was detected in response to binding soluble oviduct glycans[19]. The increased viability of sperm bound to oviduct glycans *in vitro* suggests that binding to suLe^X^ inhibits some capacitation-related events and maintains the pre-capacitation state. These changes may result in alterations in the sperm metabolic profile, allowing sperm to survive in the isthmus for longer periods. Our objective was to determine if binding to suLe^X^ altered porcine sperm metabolism in a manner that would explain the enhanced lifespan of sperm bound to oviduct glycans in the sperm reservoir.

## Results

### Metabolomic Analysis of Sperm Incubated with Oviduct Glycans

To assess the changes in sperm metabolism due to the interaction with oviduct glycans, suLe^X^ covalently attached to a 20 kDa polyacrylamide chain (referred to hereafter as suLe^X^) was added to porcine sperm and metabolite concentrations were assessed by GC/MS targeted metabolic profiling after 0.5 and 4.0 hr of incubation (Figure S1). The addition of culture medium was used as a control. For comparison, suLe^A^, an isomer of suLe^X^ that was also attached to a 20 kDa polyacrylamide chain and binds few sperm before capacitation and 20-25% of sperm after capacitation, was examined. The resulting data were submitted for quantitative enrichment analysis using the Small Molecule Pathway Database (SMPDB) metabolite database within Metaboanalyst.ca[23]. Quantitative enrichment analysis considers how many metabolites within a pathway are altered, how closely related those metabolites are to a given endpoint or each other, and the magnitude of the change between treatments to rank the pathways that are most different between the given treatments and test for statistical significance. Sperm incubated with suLe^X^ had lower concentrations of several metabolites compared with suLe^A^ or control-treated sperm at the 0.5 hr incubation time. The pathway most altered when compared to the control was ubiquinone biosynthesis, due to a significantly lower relative abundance of 4-hydroxybenzoic acid (4-HB) (Figure 1A, B). 4-HB is the immediate precursor of ubiquinone, also known as Coenzyme Q10, a co-factor in respiratory Complexes I, II, and III of the electron transport chain residing in the inner mitochondrial membrane[24-27] although it may also have roles in other cellular membranes[28]. In the electron transport chain, ubiquinone carries electrons from Complexes I and II to Complex III[29]. The ubiquinone-containing respiratory Complex II has a second role, oxidizing succinate to fumarate in the citric acid cycle, hence its alternative names succinate-coenzyme Q reductase and succinate dehydrogenase[30]. Consistent with this function, the abundance of its product fumarate but not the precursor succinate was reduced by suLe^X^ (Figure 1E). A product further downstream in the citric acid cycle, citrate, was reduced numerically although not statistically. The differential effect of suLe^X^ on fumarate and succinate further implicated ubiquinone biosynthesis as being significantly affected in sperm during the 0.5 hr incubation.

**Figure 1.**
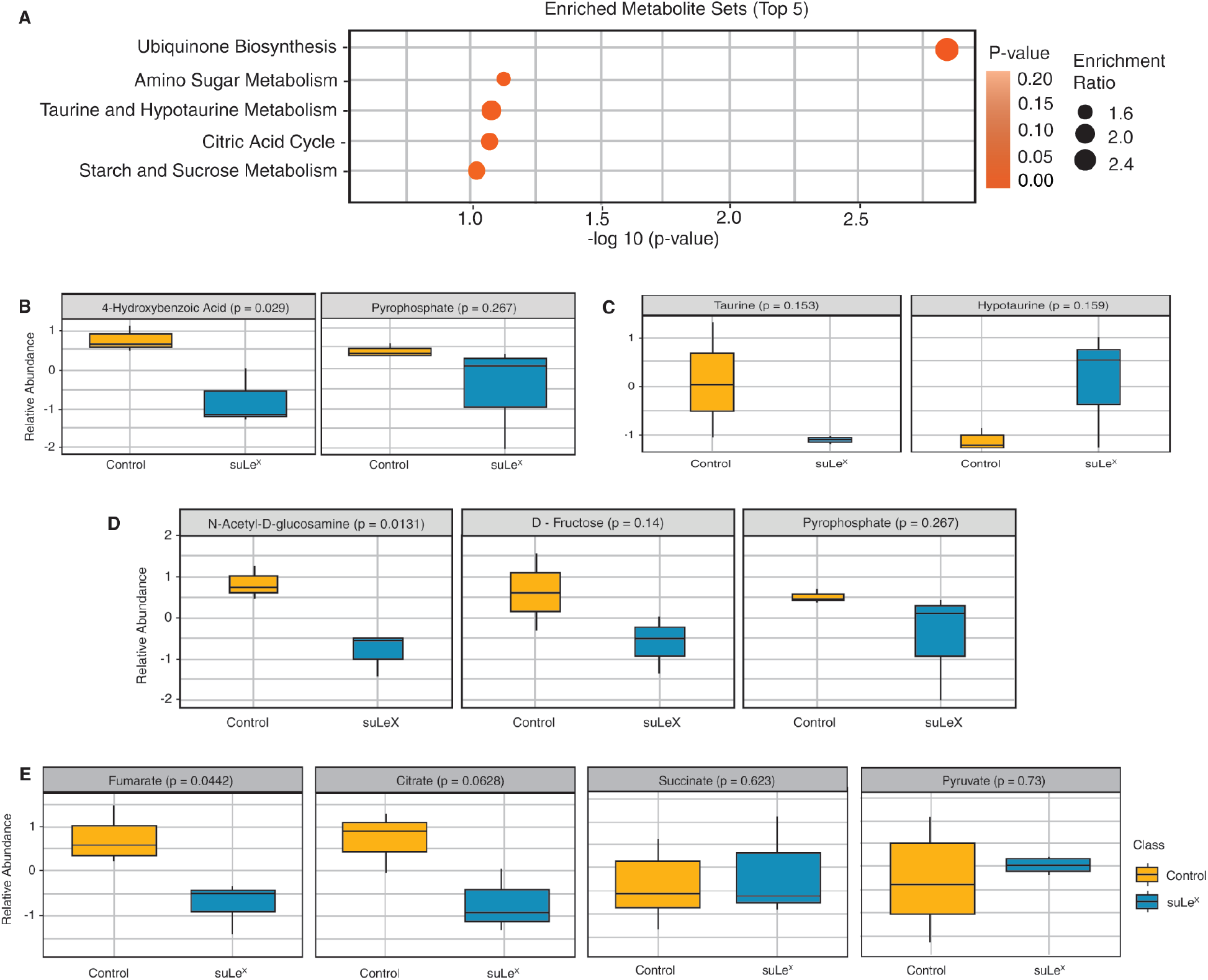
Quantitative Enrichment Analysis of Sperm Incubated with suLe^X^ Compared with Vehicle Control after 0.5 hr of Capacitation. **A**. The analysis ranked affected pathways by the magnitude of differences between the treatments and the number of altered metabolites within the pathway. The 4 most altered pathways in the treatment are listed. Ubiquinone biosynthesis was the only pathway significantly affected by suLe^X^. **B**. The precursor of ubiquinone, 4-hydroxybenzoic acid, was reduced by 76% in sperm incubated with suLe^X^. **C**. The taurine and hypotaurine metabolism pathway was implicated, but not affected significantly. **D**. Among components of the amino sugar metabolism pathway, N-acetyl-D-glucosamine was reduced but there were no other significant differences among metabolites. **E**. Among components of the citric acid cycle, fumaric acid was lower in suLe^X^ sperm and citric acid tended to be reduced but there were no changes in other components including succinate and pyruvate. Each experiment included at least 3 independent replicates. The line in the box of the box-and-whisker plots shows the overall mean, the borders of the box show the upper and lower quartile, and the ends of the lines show the 5^th^ and 95^th^ percentiles.

Additionally, the relative abundance of N-acetyl-D-glucosamine was lower in suLe^X^ sperm compared with control (Figure 1D), implicating the amino sugar metabolism pathway (Figure 1A) although the concentration of other metabolites in this pathway was not altered. Interestingly, taurine and its precursor hypotaurine were changed in opposing directions although not significantly (Figure 1C).

After 4 hrs of incubation, the time when porcine sperm have completed capacitation *in vitro*[31], the analysis of suLe^X^ compared with control sperm revealed no significant differences among pathways between the two groups (Figure 2A, B). There were also no differences noted between specific metabolites.

**Figure 2.**
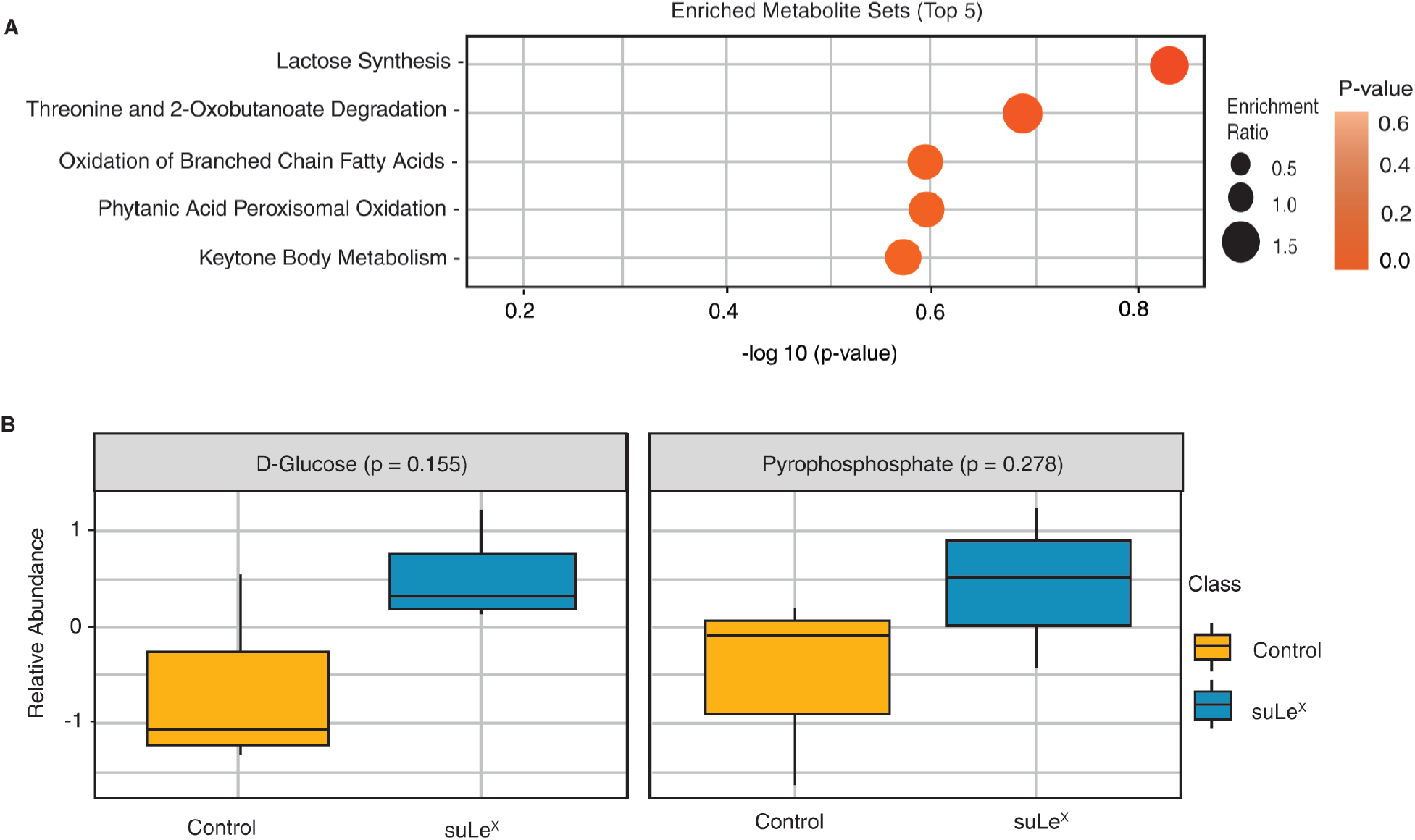
Quantitative Enrichment Analysis of Sperm Incubated with suLe^X^ Compared to Vehicle Control after 4 hr of Capacitation. **A**. The metabolic pathways most altered by exposure to suLe^X^ compared with control sperm after 4 hours of capacitation. There were no significant differences among pathways due to sperm treatment with suLe^X^. Furthermore (B), there were no differences in the major metabolites in the pathway that was most different (lactose synthesis). Each experiment included at least 3 independent replicates. The line in the box of the box-and-whisker plots shows the overall mean, the borders of the box show the upper and lower quartile and the ends of the lines show the 5^th^ and 95^t^ percentiles.

We also compared the metabolome of sperm incubated with suLe^X^ to sperm incubated with the isomer suLe^A^, a glycan that binds few sperm before capacitation but 20-25% of sperm after 4 hr of capacitation[14]. The comparison by quantitative enrichment analysis revealed that sperm incubated with suLe^X^ had significantly less cholesterol and marginally less desmosterol than suLe^A^ after 0.5 hr of incubation (Figure 3A, B). The reduction in these two steroids (Figure 3B) implicated the pathways named steroid biosynthesis and steroidogenesis (Figure 3A). Although sperm treated with suLe^X^ had less cholesterol, these pathway names are misleading for sperm because they are deficient in cholesterol and steroid biosynthesis[32]. The overall phosphatidylinositol pathway was also differentially affected (Figure 3A) although no specific metabolites of this pathway were themselves significantly different.

**Figure 3.**
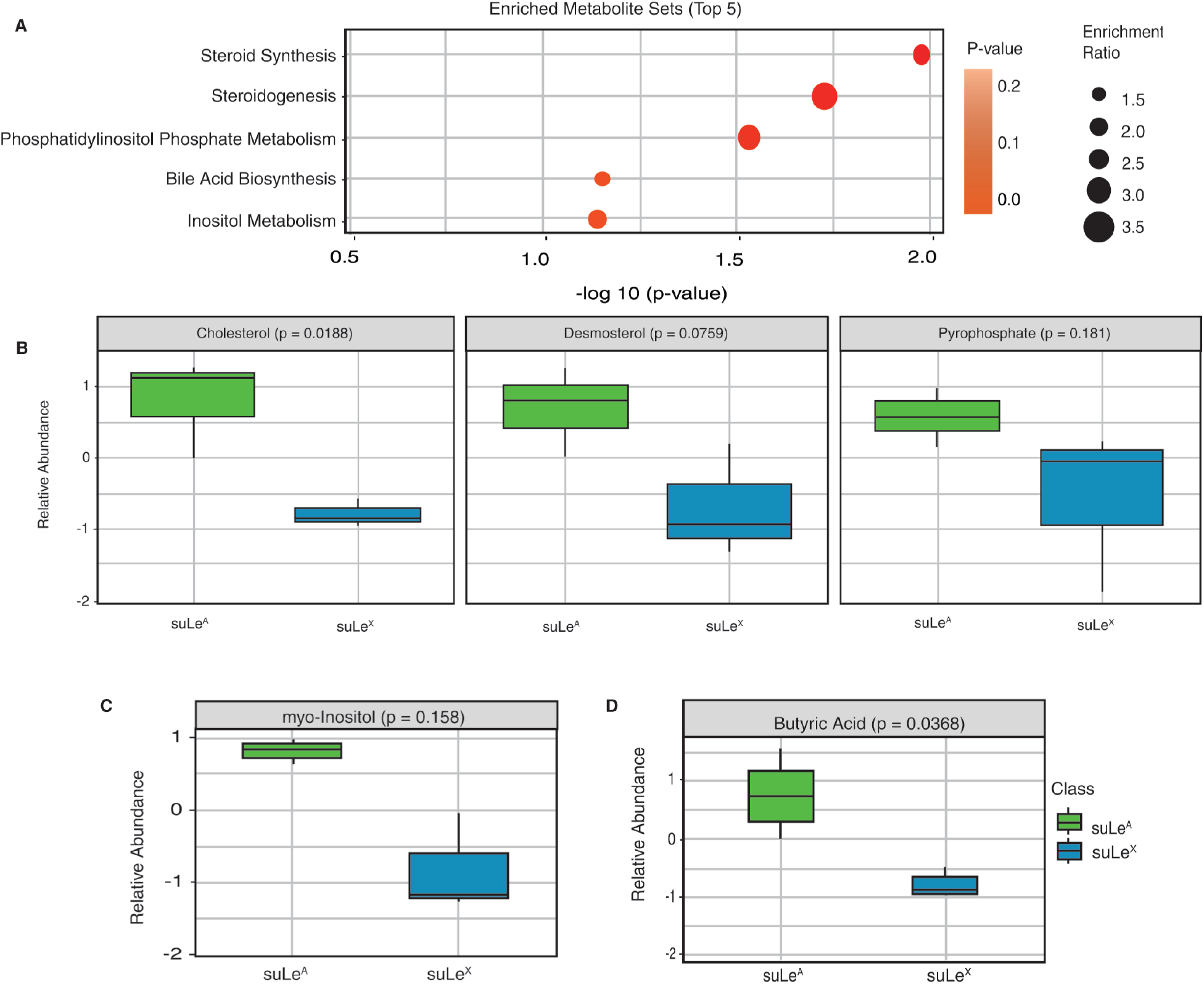
Quantitative Enrichment Analysis of Sperm Incubated with suLe^X^ Compared to suLe^A^ after 0.5 hr of Capacitation. **A**. The steroid biosynthesis and steroidogenesis pathways and the phosphatidylinositol phosphate pathway were the only pathways significantly different among both Lewis trisaccharides. The top 5 pathways are listed. **B**. Among the steroid biosynthesis and steroidogenesis pathway components, cholesterol was lower in suLe^X^-treated sperm compared with suLe^A^-treated sperm and desmosterol tended to be lower. **C**. No single phosphatidylinositol phosphate metabolism pathway component was altered at 0.5 hr but myo-inositol had the smallest P value. **D**. The bile acid biosynthesis pathway was modestly altered. Sperm incubated with suLe^X^ had less butyric acid than suLe^A^-treated sperm. Each experiment included at least 3 independent replicates. The line in the box of the box-and-whisker plots shows the overall mean, the borders of the box show the upper and lower quartile, and the ends of the lines show the 5^th^ and 95^t^ percentiles.

Comparing suLe^X^ to suLe^A^-treated sperm after 4 hours of capacitation, there was a tendency for higher cholesterol in suLe^X^-treated sperm compared with suLe^A^-treated sperm and the steroidogenesis pathway was the most differentially impacted between the two Lewis trisaccharides (Figure 4). No other pathways were significantly altered.

**Figure 4.**
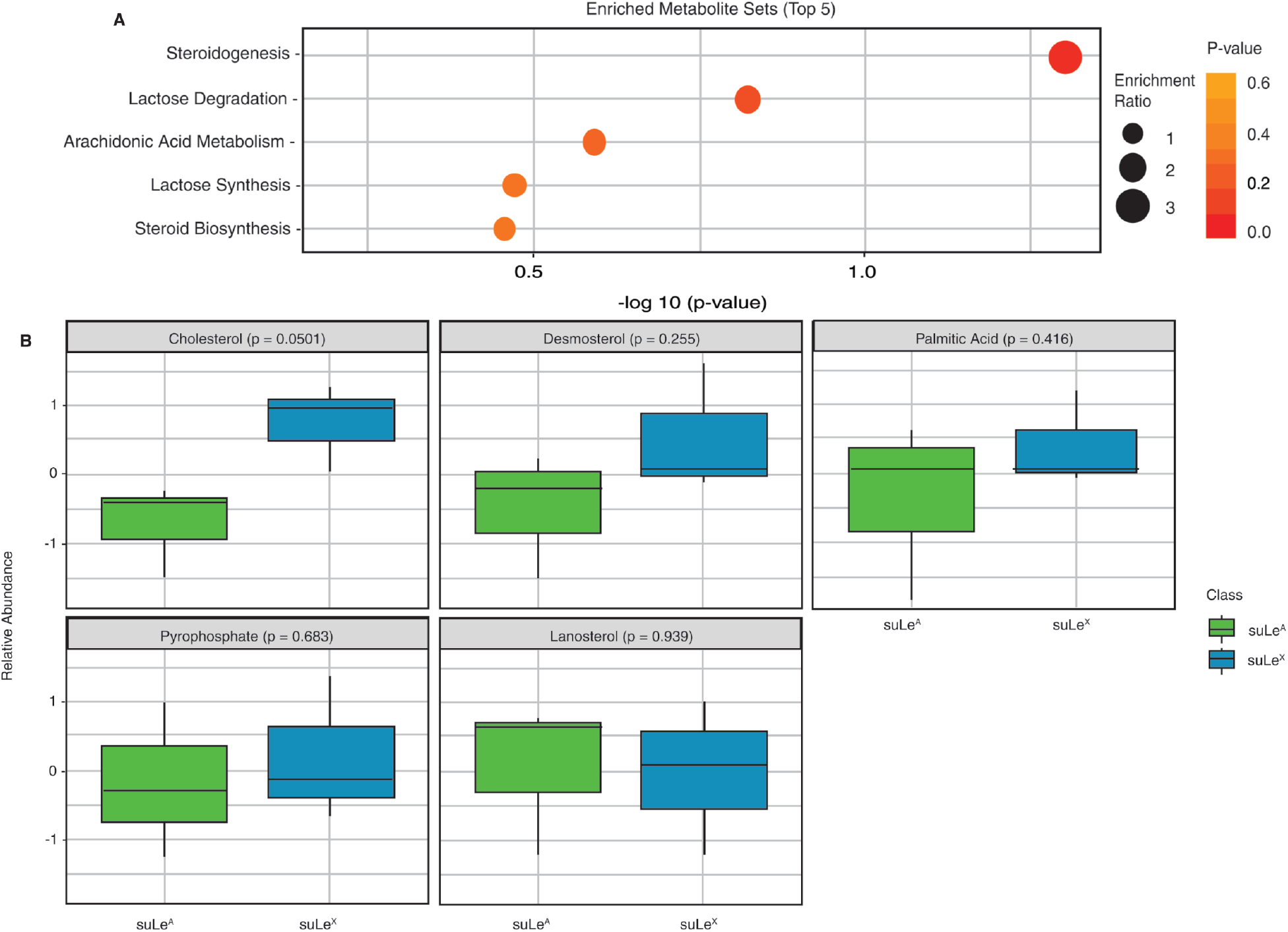
Quantitative Enrichment Analysis of Sperm Incubated with suLe^X^ Compared to suLe^A^ after 4.0 hr of Capacitation. **A**. The top 5 pathways are shown and the only significantly altered pathway was steroidogenesis. **B**. Among steroidogenesis metabolites, there was significantly more cholesterol after 4 hours of capacitation compared with sperm treated with suLe^A^. The remainder of metabolites in this pathway were not significantly different. Each experiment included at least 3 independent replicates. The line in the box of the box-and-whisker plots shows the overall mean, the borders of the box show the upper and lower quartile and the ends of the lines show the 5^th^ and 95^t^ percentiles.

Additional analyses were performed comparing suLe^A^-treated sperm and controls at 0.5 and 4 hrs of capacitation. The only significant difference was at 4 hr of capacitation; the steroidogenesis pathway was affected, due to a decrease in cholesterol in the sperm incubated with suLe^A^ (Figures S2, S3).

### Flow Cytometry Analysis of Sperm Viability and Reactive Oxygen Species (ROS)

During oxidative phosphorylation, a small proportion of electrons escape from the ETC and react with molecular oxygen to produce superoxide and oxygen radical products of superoxide such as peroxide[33, 34]. Normally, cells produce ROS in a controlled process, but excessive ROS production becomes pathological. ROS can cause DNA damage, protein oxidation, and lipid peroxidation, leading to cell death[33]. Sperm are believed to be very susceptible to ROS due to their limited ability to reduce ROS and repair DNA and membrane damage[35-38]. Oxidative phosphorylation increases during sperm capacitation[39], likely increasing mitochondrial ETC production of ROS and the potential for sperm injury.

Because ubiquinone is a component of the electron transport chain (ETC), we investigated whether suLe^X^ influences sperm viability by reducing the production of ROS. We reasoned that an outcome of the reduced ubiquinone synthesis and oxidative phosphorylation induced by suLe^X^ might be suppressed formation of ROS, which may contribute to a prolonged sperm lifespan. We assessed sperm incubated for various times under capacitating conditions using a ROS probe (CellROX Deep Red) and flow cytometry to measure total ROS quantitatively[40].

Sperm incubated with suLe^X^ had lower numbers of single events (fewer sperm not attached to other sperm) compared with sperm treated with suLe^A^ or with medium (Figure 5A). This was consistent across all times observed during capacitation. Sulfated Le^X^ also reduced the number of single live sperm at 1 hr and tended to reduce the number of single live sperm at 0.5 and 4 hr (Figure 5B). This may be explained by the tendency of more live sperm incubated with suLe^X^ to form aggregates that are gated out of the flow cytometry analysis because they are not single cells. Importantly, the overall CellROX staining intensity in single live sperm was reduced when sperm were treated with suLe^X^ at 4 hr compared with control, indicating that suLe^X^ reduced ROS accumulation (Figure 5C). There were no differences in ROS intensity at the earlier time points.

**Figure 5.**
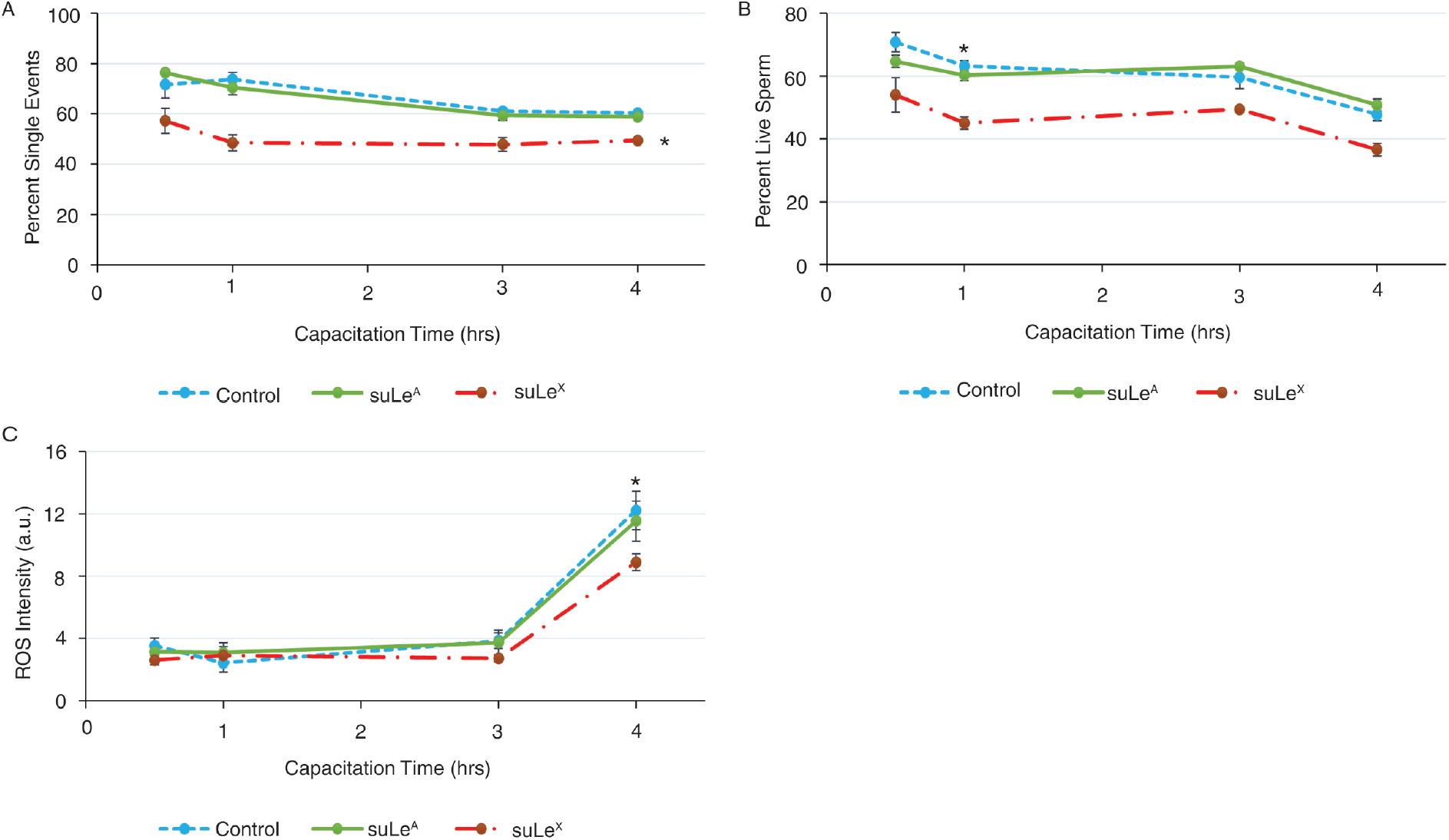
Sperm aggregation, viability, and ROS intensity after treatment with suLe^X^ and suLe^X^. **A**. SuLe^X^ reduced the percentage of single sperm events. The number of events that were defined as single sperm, based on size and density, was significantly reduced when sperm were incubated with suLe^X^ compared with other treatments throughout capacitation. **B**. suLe^X^-treated sperm had reduced viability during capacitation. The number of events that were within the threshold for both single and live single sperm was statistically reduced in suLe^X^-treated sperm compared with all other treatments at 1 hr. **C**. Mean ROS intensity of suLe^X^ treated sperm was reduced at 4 hr of capacitation. The average intensity of CellROX staining among the single live sperm was reduced in sperm treated with suLe^X^ compared with other treatments at 4 hr of capacitation. Each experiment included 6 independent replicates. The graphs show the mean and standard errors.

### Motility and Acrosome Responsiveness of Sperm

Sperm metabolism is a major contributor to ROS production[41]. Sperm motility is also related to metabolism[39] so we next examined if the sperm motility pattern was altered by incubation with suLe^X^. Sperm tail motion was assessed by CASA analysis, which infers differences in the tail beat pattern by assessing the movement of the head. This analysis did not detect any differences in motility patterns between suLe^X^, suLe^A,^ or control sperm at all time points assessed (Figure 6A-C). We also made an initial examination of whether sperm capacitation was affected by incubation with suLe^X^ or suLe^A^. To assess capacitation, we added a relatively low concentration (2 µM) of a Ca^2+^ ionophore, A23187, which induces acrosomal exocytosis in a portion of capacitated sperm within 15 min[19]. Sperm treated with suLe^X^ or suLe^A^ did not differ in their responsiveness to 2 µM ionophore stimulation after 4 hrs of capacitation, as measured by acrosomal exocytosis (Figure 6D).

**Figure 6.**
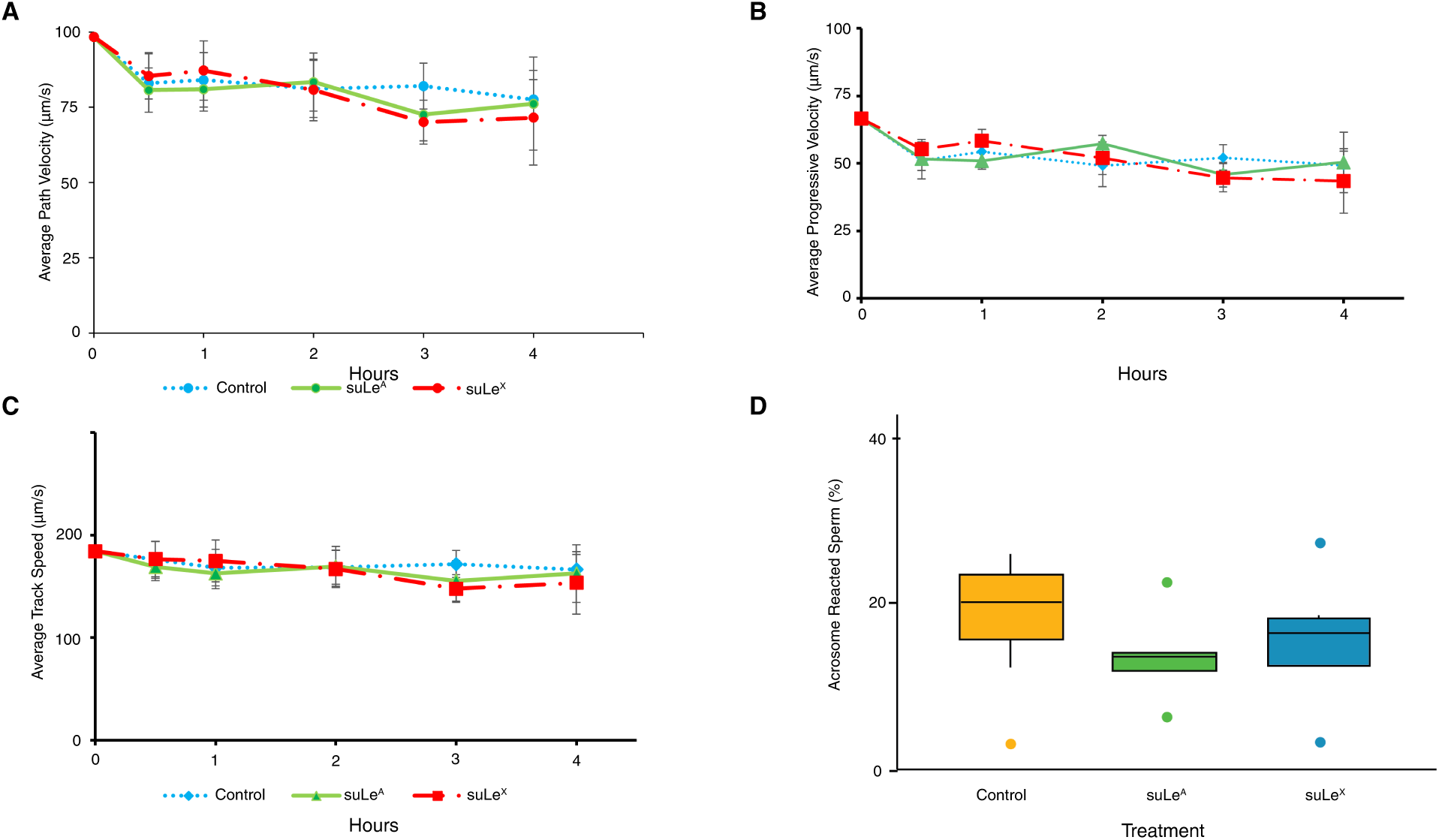
Motility and Ionophore-Responsiveness were not Altered by Treatment with suLe^X^. **A-C**. Sperm treated with suLe^X^ and suLe^A^ did not have altered motility patterns with CASA analysis. **D**. Ca^2+^ ionophore response was used as an assessment of capacitation. There were no changes in the A23187-induced acrosome reaction frequency due to suLe^X^ and suLe^A^ addition. Motility was evaluated in 3 independent replicates and acrosomal exocytosis experiments were performed 6 times. The graphs show the mean and standard errors. In **D**, the line in the box of the box-and-whisker plots shows the overall mean, the borders of the box show the upper and lower quartile, the ends of the lines show the 5^th^ and 95^t^ percentiles, and the dots show outliers.

## Discussion

Previous work established that oviduct cells suppress sperm intracellular Ca^2+^, lower membrane ROS and increase sperm longevity to form a reservoir for sperm storage[42-45]. Porcine sperm are retained in the reservoir by adhesion to oviduct cell glycans containing one of two specific motifs, either the suLe^X^ trisaccharide or a branched 6-sialylated structure[14, 15]. Both glycans are individually sufficient to suppress the capacitation-induced increase in intracellular Ca^2+^ and lengthen sperm lifespan[19] but the mechanism by which sperm survival is lengthened is uncertain. Using targeted metabolomics, we report that binding to the oviduct glycan suLe^X^ diminishes ubiquinone synthesis and suppresses the formation of specific citric acid cycle components, diminishing the activity of the electron transport chain and reducing the production of ROS. Lowered ROS production may contribute to enhanced sperm lifespan as many reports have demonstrated the toxic effects of high ROS concentrations on sperm[38, 46, 47].

The most substantial change in sperm metabolism induced by the oviduct sperm binding glycan motif suLe^X^ was the reduction of the ubiquinone precursor, 4-hydroxybenzoic acid (4-HB) at 30 min of capacitation (Figure 1B). 4-HB is the direct precursor of ubiquinone, which was not among the metabolites in the targeted metabolomics panel. The ubiquinone biosynthesis pathway is of great interest in broad sectors of study, including male reproduction[48], due to the vital nature of the end product, ubiquinone, also known as Coenzyme Q10[24-27]. Ubiquinone functions as an electron acceptor in the electron transport chain (ETC) for complexes I, II, and III and transfers the electrons to cytochrome c[29]. Ubiquinone is a lipophilic molecule that resides in the membrane of the inner mitochondrial membrane (IMM) and may be transported to other membranes within the cell[28]. Most steps of ubiquinone synthesis are carried out from the initial substrate, tyrosine, by a super-complex of proteins termed the CoQ synthome[24, 26, 49]. It is hypothesized that high concentrations of 4-HB activate its transfer into the IMM[29]. Once within the IMM, 4-HB is irreversibly prenylated by the enzyme *trans*-prenyltransferase[27]. However, the mechanisms that regulate ubiquinone synthesis and degradation are not clear.

To our knowledge, this work is the first to demonstrate that 4-HB and ubiquinone concentration are regulated in sperm. Although the half-life of ubiquinone in sperm is unknown, the catabolism of ubiquinone has been measured in the testis and its half-life is 50 hr, among the lowest of the tissues examined[50]. But this half-life is still relatively long compared to the rapid effects on sperm ROS and sperm longevity that are observed within 30 min of suLe^X^ addition. suLe^X^ most likely increases the breakdown of 4-HB and perhaps ubiquinone to reduce their concentration. Because 4-HB concentrations are decreased within 30 min after the addition of suLe^X^, degradation of 4-HB must occur swiftly. Additionally, the change in 4-HB abundance may also be due to decreased synthesis, which would also be rapid. The half-life of 4-HB in sperm is unknown. Due to the short time and very limited transcription ability of sperm, the reduction in 4-HB and ubiquinone would likely be due to post-translational modification of enzymes involved in synthesis or degradation. Changes in phosphorylation of sperm proteins are hallmarks of sperm capacitation[20] and are one possible ubiquinone synthesis and degradation regulatory mechanism.

The decrease in 4-HB and ubiquinone induced by suLe^X^ also affected the citric acid cycle. suLe^X^ reduced the concentration of fumarate after 30 min (Figure 1E). Fumarate is produced by the action of succinate-coenzyme Q reductase (the same enzyme is known as succinate dehydrogenase or respiratory Complex II). Succinate-coenzyme Q reductase employs ubiquinone as a co-factor and is part of both the citric acid cycle and the electron transport chain, with distinct actions in each[51]. Reduced activity of this enzyme would reduce fumarate concentration without affecting succinate, which is what was observed (Figure 1). Although not significantly different, citrate concentration was also numerically reduced by suLe^X^ (P=0.06). In total, this indicates that, in addition to affecting respiratory complex enzymes in the ETC, the reduction in 4-HB and presumably ubiquinone affected the citric acid cycle.

Because physiological concentrations of ROS are important for cellular signaling during capacitation, reducing ROS may delay some components of capacitation[35, 38]. A delay in capacitation may be a second way that sperm lifespan is enhanced by suLe^X^. Although we found no evidence of a delay in capacitation by assessing the ability of sperm to respond to Ca^2+^ ionophore with acrosome reactions, capacitation is a complex multi-step process. There are no single methods to evaluate in total the many processes included during capacitation in a heterogeneous population of sperm. Furthermore, a limited number of capacitation processes may be influenced by diminished ROS production. But it was interesting that, when sperm treated with suLe^A^ were compared to sperm incubated with its isomer, suLe^X^ at 4 hr when capacitation was completed, more cholesterol was found in sperm treated with suLe^X^ (Figure 4A, B). This is consistent with the hypothesis that suLe^X^ delays some aspects of capacitation. Curiously, at 0.5 hr, sperm incubated with suLe^X^ contained less cholesterol than sperm with suLe^A^ (Figure 4). The gain in cholesterol in response to suLe^X^ at 4 hr may be due to reduced ROS production observed in sperm treated with suLe^X^ (Figure 5). There is evidence that ROS may oxidize cholesterol making it more hydrophilic[22, 52, 53] which may promote cholesterol release from sperm. Therefore, a reduction in sperm ROS may quell cholesterol release during capacitation.

Total ROS content was measured in single live sperm using flow cytometry with CellROX, a ROS fluorescent probe. Unexpectedly, the suLe^X^-treated sperm had significantly fewer single sperm compared with controls and suLe^A^. This may be explained by the bridging of sperm by suLe^X^ which, in these experiments, was attached to a 20 kDa polyacrylamide chain. When suLe^X^ was included in the capacitating medium, many of the sperm formed clusters. The clusters of agglutinated sperm are difficult to evaluate but likely included more live than moribund sperm because of the high affinity of live uncapacitated sperm for suLe^X^ compared to capacitated or dead cells[14]. The live uncapacitated sperm that bind suLe^X^ are expected to be preferentially sequestered in the agglutinated population that is gated out by flow cytometry. Consequently, there were fewer live single sperm in the suLe^X^ treatments after 1 hr of incubation and smaller non-significant differences at other times.

Before 4 hr of incubation, the mean ROS concentration in single-live sperm was not different between sperm incubated with the two Lewis structures and medium alone. But at 4 hr of capacitation, the ROS concentration in sperm incubated in medium increased markedly. The increase was blunted by suLe^X^ but not suLe^A^. No significant difference in ROS due to suLe^X^ was observed before 4 hr so there was a delay between suppressed ubiquitin synthesis and reduced ROS accumulation. This may be due to the time required for the catabolism of 4-HB and ubiquinone. The effects of mitochondrial ubiquinone abundance on sperm ROS production are in agreement with reports in somatic cells showing that greater ubiquinone concentration is associated with higher ROS production[54].

If sperm had lowered ETC activity due to reduced ubiquinone, the reduction in oxidative phosphorylation could affect sperm motility and capacitation. But there was no effect on motility detected using CASA analysis, consistent with our prior findings[19]. We assessed capacitation by the frequency of induced acrosome reactions using conditions in which sperm that have not completed capacitation do not acrosome react[19]. Although there is evidence that oxidative phosphorylation increases during sperm capacitation[39], we did not detect any change in the frequency of ionophore-stimulated acrosome reactions in sperm incubated with suLe^X^ after 4 hr. Thus, there seems to be adequate oxidative phosphorylation for some sperm functions in the presence of soluble suLe^X^.

Previous experiments testing the effects of glycans on sperm viability have often used glycans immobilized on 30-40 μm beads [14, 16, 19] rather than soluble free trisaccharides or trisaccharides attached to a 20 kDa polyacrylamide chain to resemble the multiple copies of glycans found frequently on a glycoprotein. The immobilized glycans are likely better mimics of glycans attached to glycoproteins and glycolipids on oviduct cells. But soluble glycans affect sperm behavior, such as suppressing the increase in intracellular Ca^2+^ that normally occurs during capacitation [19]. Due to the difficulty of using glycans on 30-40 μm beads for flow cytometry, experiments herein used only glycans attached to a soluble polyacrylamide chain. But immobilized glycans may have a more significant effect on sperm metabolism than soluble suLe^X^.

Very recently, it was demonstrated that oocytes lengthen their lifespan by eliminating Complex I of the ETC to also minimize ROS production and increase lifespan[55]. Although the lifespan of primordial oocytes is much longer than sperm, it is interesting that both gametes reduce ETC activity and ROS production to lengthen the time they are viable. Mammalian sperm have a smaller reduction in ETC activity, possibly because ROS have physiological roles in capacitation and the acrosome reaction that must be fulfilled. Inhibition of ubiquinone biosynthesis may also have a role in organisms. Perturbation of the *ckd1*/*coq7* gene that encodes a ubiquinone synthesis enzyme reduces oxidative stress and increases longevity in both *c. elegans* and mice[25, 56, 57].

In summary, targeted metabolomic studies showed that the oviduct glycan motif, suLe^X^, which lengthens sperm lifespan, reduced the abundance of sperm 4-HB, a precursor of ubiquinone after 30 min of incubation in capacitating medium. Along with the reduction in ubiquinone, a component of Complexes I, II, and III in the electron transport chain. ROS accumulation in sperm was reduced at 4 hr of incubation. Complex II is also known as succinate-coenzyme Q reductase for its role as a critical enzyme in the citric acid cycle. The activity of this ubiquinone-containing enzyme was reduced, as there was less of the enzyme’s product, fumarate. Thus, the citric acid cycle and oxidative phosphorylation were both suppressed by suLe^X^. Reduced ROS at the end of capacitation in sperm treated by suLe^X^ did not affect sperm motility or the ionophore-induced acrosome reaction but was associated with cholesterol loss. Reduced ubiquinone synthesis muted ROS’s pathological effects, lengthening sperm lifespan. These data provide a mechanistic understanding of how an oviduct component, the glycan suLe^X^, lengthens sperm lifespan and are the first to implicate the ubiquinone synthesis pathway in sperm function. The results are also broadly applicable as an example of a rapid novel effect that a cell matrix glycan can have on cellular metabolism.

## Materials and Methods

Porcine semen was a gift from PIC North America. Semen was collected by applying manual pressure to the glans penis. Semen was extended in BSA-free NeutriXcell (IMV, Maple Grove, MN), cooled to 17° C, shipped to the laboratory, and processed within 24 hr. 3-O-sulfated Lewis X and 3-O-sulfated Lewis A were coupled to a 20-kDa polyacrylamide chain resulting in a 0.8-0.9 µg trisaccharide per mg polyacrylamide conjugate ratio as described [58, 59] and will be referred to herein as suLe^X^ and suLe^A^ respectively. Reagents used in flow cytometry analysis were CellROX Deep Red (Thermo Fisher, Waltham, MA), propidium iodide (Molecular Probes, Eugene, OR), and calcium ionophore A23187 (Millipore, Burlington, MA).

### Sperm Preparation

Extended semen (30 ml for flow cytometry, motility assays, or 200 ml for metabolomics) was centrifuged for 10 min at 600 x g, resuspended in 10 ml of dmTAPG (2 mM CaCl_2_, 4.8 mM KCl, 95 mM NaCl, 1.2 mM KH_2_PO_4_, 15 mM HEPES, 25 mM NaHCO_3_, 5.56 mM glucose, 0.6% bovine serum albumin (Fraction V, 95% purity, Sigma Aldrich, St. Louis, MO), 1 mM sodium pyruvate, and centrifuged for an additional 5 min at 400 x g. Sperm were then resuspended in dmTAPG at a concentration of 10^7^ sperm/ml supplemented with one of the following: 0.01 µg/ml 3-O’-sulfated-Lewis X (suLe^X^), 0.01 µg/ml, 3-O’-sulfated-Lewis A (su-Le^A^), or vehicle control, dmTAPG. An outline of sampling times is shown in Figure 1.

### Metabolomic Analysis

Metabolomic analysis samples were prepared according to the above protocol for sperm preparation and incubated with suLe^X^, suLe^A,^ or control (medium only) for 0.5 and 4 hours at 39 °C under capacitating conditions in dmTAPG. At the end of capacitation, sperm were centrifuged at 3,000 x g for 5 min to remove the treatment medium. The sperm pellet was resuspended in Hanks Balanced Salt Solution (HBS, Gemini Bio, Sacramento, CA), and samples were centrifuged for a total of 3 washes in HBS to remove the treatment medium. The sperm pellet was stored at -80° C until analysis. This experiment was repeated across three independent replicates using semen pooled from 2-3 different groups of boars for each replicate.

Metabolic sample preparation and targeted GC/MS metabolomic profiling were performed by the Metabolomics Center, Roy J. Carver Biotechnology Center, University of Illinois at Urbana-Champaign using standard protocols [60].

### Flow Cytometry

Sperm were incubated with the polyacrylamide-conjugate of suLe^X^, or suLe^A^ or control (medium only) under capacitating conditions for 0.5, 1, 3, and 4 hours at 39° C at 10 × 10^5^ cells/ml in 4 ml. Thirty min before analysis, sperm in all treatments were stained with CellROX Deep Red at a final concentration of 2.5 µM and returned to 39 °C. Samples were transported to the flow cytometry lab (10 min with warm packs at 39 °C) and incubated at 37 °C for the duration of the experiments. Five min before initial flow cytometry data collection, samples were stained with 0.1 µg/ml propidium iodide (PI). Flow cytometry analysis was performed using a Thermo Fisher BD II flow cytometer at the University of Illinois Urbana-Champaign Cytometry and Microscopy to Omics Core within the Roy J. Carver Biotechnology Center. Sperm from each treatment were analyzed for viability, using PI, and the presence of ROS, using CellROX Deep Red at 0.5, 1, 3, or 4 hours of capacitation.

Flow cytometry data were sorted via the following gating strategy (Figure S4). Total events fell into the typical flame shape when side scatter (sperm complexity, relating to laser scattering due to intracellular structures) was graphed against forward scatter (sperm size) properties. The population selected from total events was the most uniform in size and assumed to be single cells. Within this single-cell population, the events were gated by propidium iodide staining intensity, which fell into two distinct populations. The low PI-staining population was accepted as live sperm. Ten thousand single, live sperm were measured for ROS intensity at each capacitation time. Live cells were gated into a low or high CellROX staining population corresponding to low and high ROS.

Samples were incubated with 2 µM calcium ionophore (A23187, final concentration) in dmTAPG to induce the acrosome reaction. Subsequent flow cytometry analysis was performed 15 min after ionophore addition. The resulting flow cytometry data were analyzed using FCS Express v.6 (Denovo Software, Los Angeles, CA).

### Motility Analysis

At 0.5, 1, 2, 3, and 4 hours of capacitation in the treatment medium, sperm motility was measured by computer-assisted sperm analysis (CASA, Hamilton Thorne, Beverly, MA). A volume of 4 µL of each sample was collected and drawn onto a disposable Leja 4 chamber (slide of depth 19.7 µm) and placed on a 37° C prewarmed stage. Each sample was evaluated by assessing at least 1,000 sperm from 10 randomly selected fields. Using the default parameters in the porcine module, the percentage of total motility, the percentage of progressive motility, average path velocity (VAP, µm/s), progressive velocity (VSL, µm/s), and track speed (VCL, µm/s) were recorded.

### Acrosome Reaction Assessment

During the flow cytometry experiments, a 100 µl aliquot of sperm from each sample at each time point was collected and incubated for 15 min with 2 µM calcium ionophore A23187 or vehicle control, fixed in 4% formaldehyde, washed with 0.1% ammonium acetate, spread on a slide, air-dried, and stained with Coomassie Blue G-250 [61]. The percentage of sperm that had undergone an acrosome reaction was evaluated in at least 200 sperm per sample. Each sample was counted by two independent readers and counts were averaged between readers. If the first two counts were more than 10 % different, the count was independently repeated by both readers until the counts were within 10 %.

### Statistical Analysis

Metabolomic analysis was carried out using the Metaboanalyst.ca platform, version 5.0 [23]. Quantitative enrichment analysis was performed on samples within each time point compared with the control. Data were normalized by dividing each variable by the overall median of each metabolite within a group. These values were log-transformed and scaled (mean-centered and divided by the standard deviation of each variable). The library selected was the SMPDB Metabolic pathway associated metabolite set, containing 99 entries.

Flow cytometry data were analyzed using R with the agricolae, DescTools, and stats packages. The endpoints analyzed were the percentage of sperm that had undergone the acrosome reaction, the percentage of sperm within the single cell gate, the percentage of sperm within the live threshold gate (low PI), the percentage of live sperm within the low and high ROS gates and the median intensity of CellROX staining within the live population, normalized to the number of single-live events. This normalized intensity did not follow a normal distribution so the square root transformation and scaling by a factor of 1,000 was applied for analysis. The ANOVA procedure was used to test for main effects followed by a two-tailed Tukey HSD post hoc analysis. Significance was determined by a p-value of ≤ 0.05. Data are presented as untransformed arbitrary units and percentages, means, and standard errors.

## Non-Standard Abbreviations

suLeX: 3-O-sulfated Lewis X trisaccharide
bi-SiaLN: Biantennary 6-sialylated oligosaccharide

## Acknowledgements

Research reported in this publication was supported by the Eunice Kennedy Shriver National Institute of Child Health and Human Development of the National Institutes of Health under award number RO1HD095841. The content is solely the responsibility of the authors and does not necessarily represent the official views of the National Institutes of Health. The authors thank Brett Nixon for insightful comments on the manuscript.

## Author Contributions

JH and DM conceived the project and designed the experiments, JH and KM performed the experiments, JM and DM interpreted the results, NB provided reagents and advice in their use, JH and DM wrote the manuscript and JH, NB, and DM edited the manuscript.

