## Supplemental Figures for "An oviduct glycan increases sperm lifespan by diminishing ubiquinone and production of reactive oxygen species"

**Supplementary Figures**

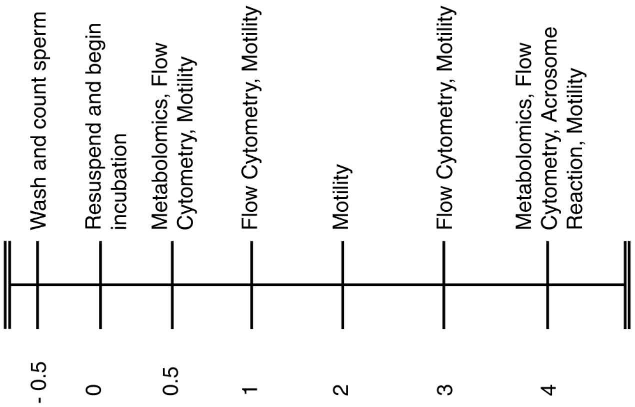


**Figure S1: Schematic of Experimental Timeline.** The time of specific observations is represented as hours of sperm incubation. The analyses that were performed are indicated.

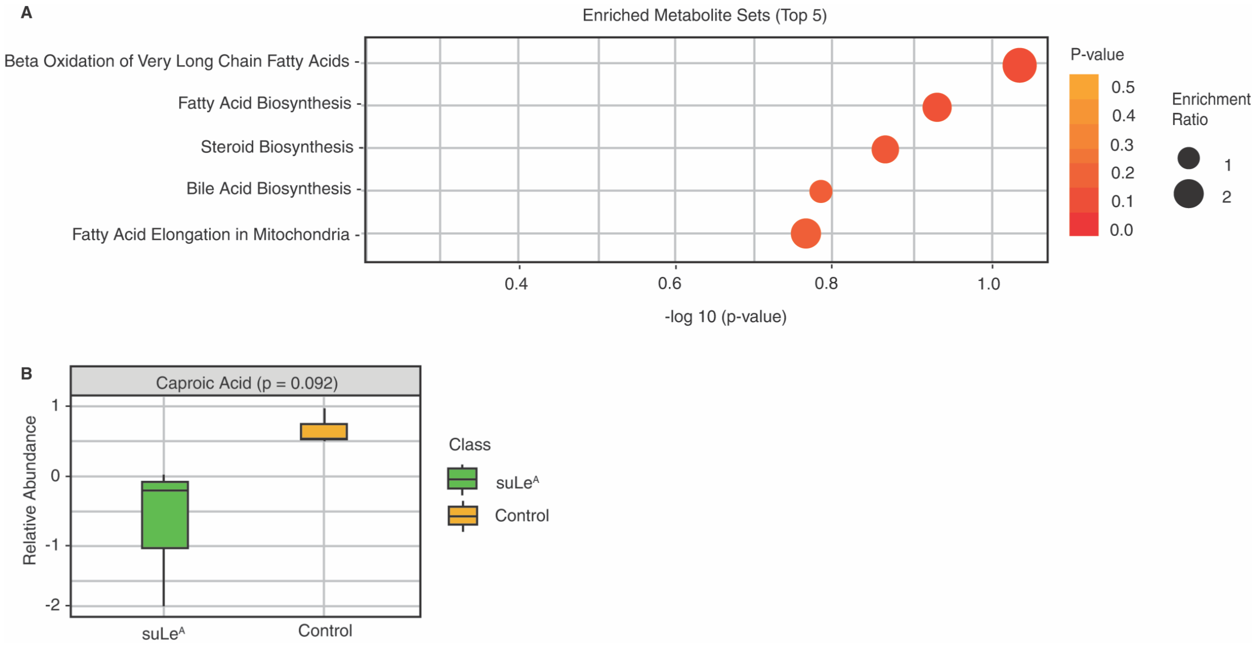


**Figure S2. Quantitative Enrichment Analysis of Sperm Incubated with suLe$^{A}$ Compared with Vehicle Control after 0.5 hr of Capacitation. A.** The top 5 pathways are shown. The pathway that differs the most between suLe$^{A}$ and control was beta-oxidation of long-chain fatty acids which was only marginally significant. **B.** Caproic acid tended to be lower in sperm incubated with suLe$^{A}$ compared with vehicle control after 0.5 hrs of incubation. The abundance of other metabolites was not changed.

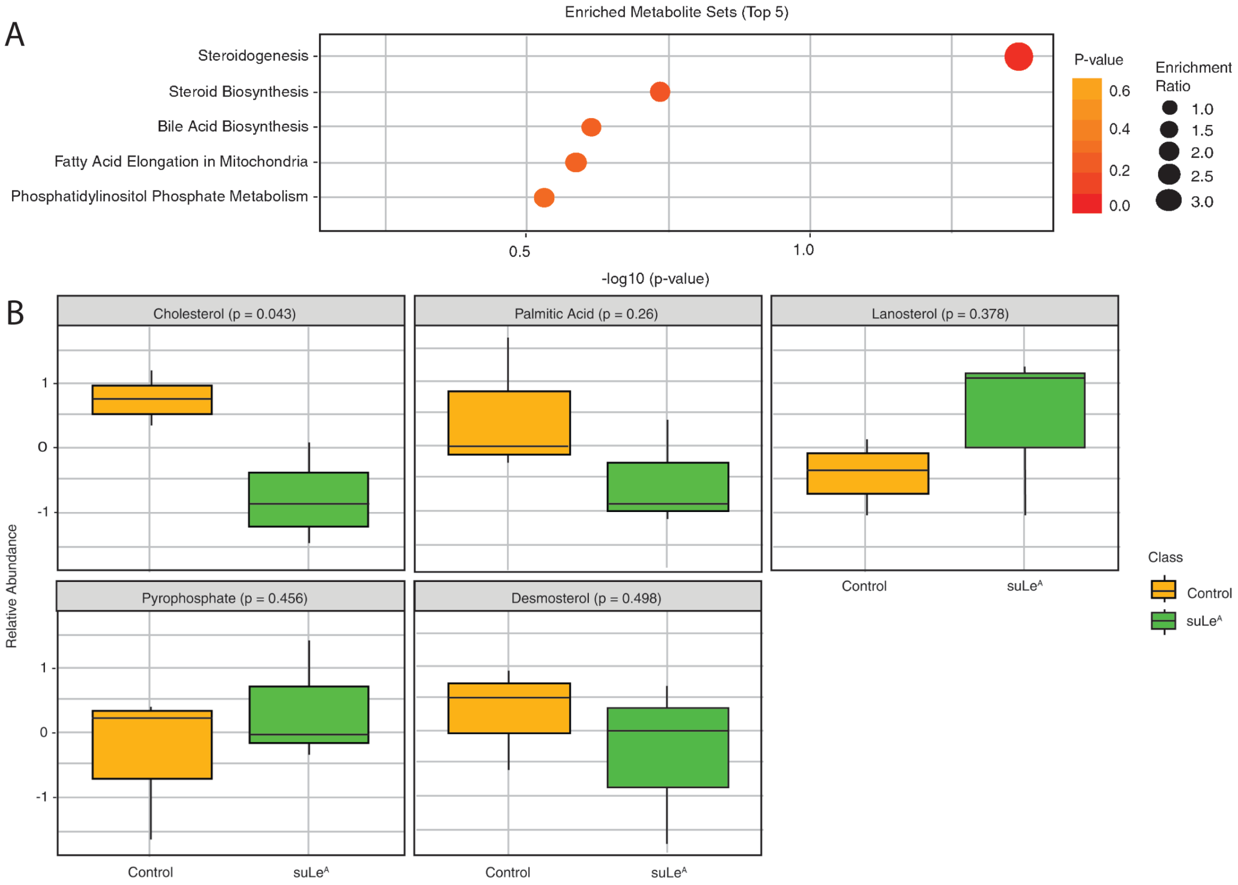


**Figure S3. Quantitative Enrichment Analysis of Sperm Incubated with suLe$^{A}$ Compared with Vehicle Control after 4.0 hr of Capacitation. A.** The top 5 pathways are shown. The only pathway that was influenced by suLe$^{A}$ compared to vehicle control was the steroidogenesis pathway. **B.** Only cholesterol was altered, and it was reduced compared to the control medium.

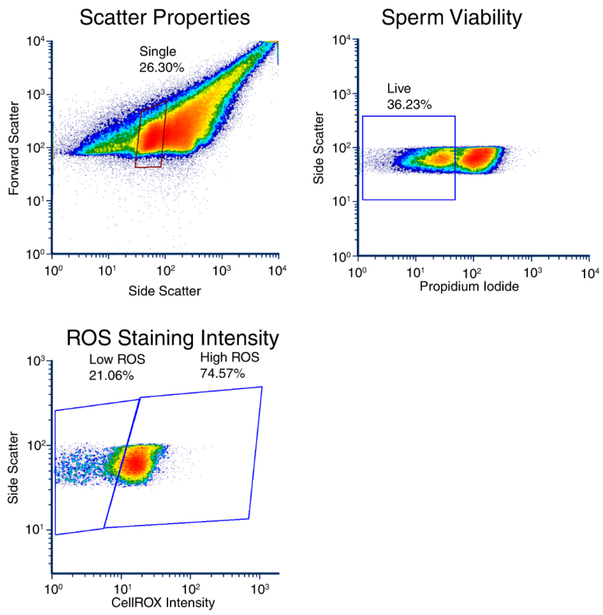


**Figure S4 Measurement of ROS in single live sperm by flow cytometry. A.** Events within each sample were first sorted by size (side scatter) and density (forward scatter). The blue box indicates events selected as the most uniform in size. This subpopulation of sperm was gated for subsequent analysis. **B.** The subpopulation selected in A was gated for propidium iodide staining intensity (viability). The events falling within the blue box of B were selected as live sperm. **C.** Live sperm were then assessed using CellROX Deep Red staining. The population in the left box was designated low staining (low ROS) and the right box was designated high staining.
